# ECCFP: a consecutive full pass based bioinformatic analysis for eccDNA identification using Nanopore sequencing data

**DOI:** 10.1101/2025.05.13.653627

**Authors:** Tangxuan Zhang, Wang Li, Qingsong Zen, Jun Zhang, Biyuan Miao, Mengting Li, Juan Luo, Tianliang Liu, Shifu Chen, Shaogui Wan

## Abstract

It is commonly known that extrachromosomal circular DNA (eccDNA) has the potential as a molecular marker because of its close relationship with cancer progress and its prevalent existence in eukaryotic organisms. The mainstream technique of eccDNA detection is using high-throughput sequencing supported by bioinformatics analysis. Although these have various analysis pipelines for sequencing data, they are restricted by sequencing platforms or have shortcomings in accuracy and efficiency. To address these limitations, we design ECCFP, a bioinformatic analysis pipeline that detects eccDNAs amplified by rolling circle amplification (RCA) from long-read sequencing data and outputs eccDNA genomic coordinates and consensus sequences. This pipeline proposes a rigorous algorithm to retain all consecutive full passes derived from individual reads to obtain candidate eccDNAs, followed by systematic consolidation of candidate eccDNAs to detect unique eccDNAs. Using simulated datasets and experimental eccDNA sequencing datasets, we estimated ECCFP in several aspects and compared it with other existing pipelines. It exhibits a marked reduction in false positive rates compared with eccDNA_RCA_nanopore and superior sensitivity relative to CReSIL and FLED. Besides, inverse PCR and Sanger sequencing further validated the existence and accuracy of the position of detected eccDNAs by ECCFP. Collectively, ECCFP provides a more efficient choice for eccDNA detection from long-read sequencing data.

## Introduction

Extrachromosomal circular DNA (eccDNA), which exists stably independent of chromosomes, is widely distributed in eukaryotes. In recent years, eccDNA has garnered increasing attention for its roles in cancer initiation, progression, intratumoral heterogeneity, and drug resistance^[1–6]^, while also demonstrating immense potential in non-invasive diagnostic applications. Since its primal observation in human cells via electron microscopy^[7]^, research on eccDNA has progressively advanced. Subsequent studies reveal that eccDNA contains tandem repeat sequences from chromosomes during Drosophila melanogaster development using two-dimensional gel electrophoresis^[8]^. With the rapid evolution of technology, ecDNA typically greater than 500 kilobases (kb) in size and containing genes^[9]^ has been identified in nearly half of the cancers and confirmed through fluorescence in situ hybridization (FISH) probes^[10,11]^. EcDNA plays a pivotal role in oncogenesis and malignant progression because it facilitates intratumoral heterogeneity. Additionally, microDNA (200–400 bp) has been shown to regulate gene expression by generating novel regulatory RNAs^[12]^, as demonstrated by dual-luciferase assays^[13]^. Direct quantification of microDNA levels via qPCR with outward-facing primers revealed that its abundance is influenced by resection after double-strand DNA breaks (DSBs) and repair via microhomology-mediated end joining (MMEJ)^[14]^. These discoveries present the biological significance of eccDNA in genomic regulation and disease pathogenesis.

Currently predominant approaches for eccDNA detection rely on high-throughput sequencing technologies integrated with bioinformatic analyses[15,16]. These approaches include whole-genome sequencing (WGS)^[17]^, assay for transposase-accessible chromatin sequencing (ATAC-seq)^[18,19]^, and eccDNA sequencing (eccDNA-seq)^[20]^. Among these, eccDNA-seq significantly enhances detection efficiency by rolling circle amplification (RCA) to enrich eccDNAs. Next-generation sequencing technology is usually used to detect eccDNA, and the corresponding bioinformatics analysis pipeline is commonly used Circle-Map^[21]^, AmpliconArchitect (AA)^[22]^ and GCAP^[23]^, that can be used to obtain the quantity and location of eccDNA molecules for further analysis. However, short-read sequencing technology faces the problem of reassembling eccDNA with complex structures, such as repeat tandem, and these analysis pipelines cannot output consensus sequences.

In particular, the Oxford Nanopore sequencing platform has been increasingly employed for eccDNA detection^[24]^, leveraging its advantages in assessing amplified DNA quality^[25,26]^. Regrettably, existing analytical pipelines for long-read sequencing data exhibit notable limitations. For instance, Ciderseq2, primarily designed for circular DNA detection in viral and plant systems, has not yet been tested for eccDNA analysis from animal cells^[27]^. Although ecc_finder is capable of processing diverse data types, it demonstrates low sensitivity in eccDNA identification from long-read sequencing datasets^[28]^. The pipeline named circular-calling imposes elevated technical demands on users, requiring proficiency in Snakemake software^[29]^. Although CReSIL and FLED provide positional information of eccDNAs, their practical applications have deficiency in accuracy of junction site determination and consensus sequence output^[30,31]^. Furthermore, while eccDNA_RCA_nanopore enables detection of eccDNA molecules in individual reads, it suffers from high false positive rates^[32]^. These limitations collectively highlight the need for further optimization of current eccDNA analysis pipelines in terms of accuracy and efficiency. To address these limitations of the current methods, we developed a new analysis pipeline - EccDNA Caller based on Consecutive Full Pass (named as ECCFP). This pipeline integrates the advantages of both full pass and fragment-based analysis pipelines by retaining all consecutive full pass from single read and consolidating candidate eccDNA regions to identify circular DNA molecules. Compared to current analysis pipelines, ECCFP retains all consecutive full passes—high-quality tandem repeat units derived from RCA—on each read in order to maximizing data utilization while minimizing false positives through a consolidating strategy. ECCFP not only significantly reduces false positives but also enables more precise determination of circular junction sites, particularly for eccDNAs whose flanking regions without microhomology sequences. Through analysis of simulated datasets and experimental eccDNA sequencing data from HepG2 cell lines, we demonstrated the superior efficiency and accuracy of ECCFP in eccDNA detection. Furthermore, the detection results were validated by PCR amplification and Sanger sequencing, confirming the reliability and practical utility of this method.

## Material and methods

### Cells cultivation and crude DNA isolation

The human hepatoma cell lines HepG2 was obtained from Procell Life Science&Technology (Cat No: CL-0103). HepG2 cell were cultured in Minimum essential medium (MEM) supplemented with 10% fetal bovine serum (FBS) and 1% penicillin-streptomycin. HepG2 cells were cultivated in dished coated with tissue culture medium and maintained at 37 °C in an atmosphere containing 5% CO2. They were routinely checked for mycoplasma contamination and authenticated based on their morphology. The HepG2 cells were harvested during the exponential growth phase.

Approximately 2×10^6 cells were washed twice with cold phosphate buffer saline (PBS) at pH7.4 . DNA extraction was then performed using the E.Z.N.A. Endo-free Plasmid Mini Kit (omega) according to the manufacturer’s instructions. The concentration of the extracted DNA was determined using a Qubit 4.0 (Thermo Fisher Scientific).

### EccDNA enrichment and RCA amplification

In order to linearize the mitochondrial DNA in each sample, approximately 10 μl of extracted HepG2 DNA was treated with the genomic rare-cutting endonuclease Quick Digest MssI (Thermo Fisher Scientific) for human samples. The reaction was incubated at 37 °C for 6 hours, followed by an incubation at 65 °C for 10 minutes to inactivate the endonuclease Fast Digest MssI . Exonuclease V (NBE) was employed to digest linear DNA at 37 °C for 6 hours, followed by an incubation at 70 °C for 10 minutes to inactivate exonuclease V. After digesting linear and mitochondrial DNA, qPCR was utilized to confirm the presence of linear chromosomal and mitochondrial DNA in the product. We used a gene absent from eccDNA as a marker to confirm that the linear DNA digested completey31. The digested eccDNA was subsequently extracted using a phenol: chloroform: isoamyl alcohol (PCI) solution (25:24:1) in a Phase Lock Gel tube (TIANGEN) to minimize DNA loss. Following precipitation with carrier glycogen (Roche) and 1/10 volume of 3 M sodium acetate (pH 5.5, Solarbio), the precipitated eccDNAs were resuspended in TE buffer(pH 7.5, Integrated DNA Technology).

EquiPhi29 DNA polymerase (Thermo Fisher Scientific) is utilized for rolling circle amplification (RCA) to ensure effiectient plasma template amplification per reaction. Each 20 μl reaction mixture contained the following components:2 μl of 10×reaction buffer (Thermo Fisher Scientific), 2 μl of 10 mM dNTPs (NEB), 1 μl of Exo-Resistant Random Primer (Thermo Fisher Scientific), 0.4 μl of 100 mM DTT, 0.4 μl of 20 ng/ul BSA, 0.6 μl of inorganic pyrophosphatase (from yeast), 1 μl of EquiPhi29 DNA Polymerase, and 10 μl of eccDNA, with nuclease-free water added to reach a maximum volume of 20 μl. The reaction mixture was incubated at 30 °C for 14 hours and incubation at 65 °C for 5 minutes. The RCA product that has the potential to block nanopore and reduce sequencing yield will be decrease. Subsequently, the RCA product was subjected to additional treatment using T7 endonuclease (NEB) at 37 °C for 15 minutes to remove debranching structures. Subsequently, the debranched DNA was purified using 0.6x SPRI select beads (Beckman Coulter). The DNA concentration was measured using the Qubit 4.0 fluorometer.

### Library preparation and sequencing

Debranched DNA was employed as input, following the manufacturer’s instructions, for constructing the sequencing library using a Ligation Sequencing Kit (Oxford Nanopore Technology [ONT], SQK-LSK109) and Native Barcode Kit (ONT). The library was sequenced on GridION instrument using a flow cell (R9.4.1, FL-MIN106D) following the manufacturer’s instructions.

### Simulated Data Generation and Evaluation Parameter Definitions

To comprehensively evaluate the performance of current analysis pipelines and ECCFP, we generated simulated datasets reflecting the characteristics of each pipeline’s output using various parameters. The simulated eccDNA sequences were designed to mimic the size and chromosomal distribution of public eccDNA sequencing data from JJN3 cells^[30]^. A Python-based in-house program was developed to generate 5,000 circular sequences (as positives) and 5,000 linear sequences (as negatives) from the hg38 reference genome according to length distribution, chromosome distribution of eccDNA in a given template(Figure 4A). The program incorporated a K-nearest neighbors (KNN) algorithm to simulate the number of rolling circle amplification (RCA) cycles, ensuring alignment with real-world data distributions. Training data from previous studies (e.g., CReSIL and eccDNA_RCA_nanopore) were used to calibrate the KNN model, ensuring consistency between simulated and empirical eccDNA length distributions.

According to the actual sequencing data, reads can be approximately categorized as five types (Figure 2A):

- Type1: complicated tandem repeat sequences formed by multiple eccDNA templates;
- Type2/type3: sequences containing a single discontinuous full pass inserted at the head (type2) or tail (type3), simulating real errors in sequencing reads;
- Type4: sequences containing multiple consecutive full pass with randomly generated non-genomic sequences (20-240 bp) inserted, simulating amplification artifacts;
- Type5: sequences containing consecutive full passes.

Simulated reads containing ghost sequences were generated to approximate the complexity of actual sequencing data (Figure 4B). PBSIM3 software was employed to generate long-read sequencing data with parameters^[33]^: --strategy wgs --method qshmm --qshmm QSHMM-ONT-HQ.model --accuracy-mean 0.95 --accuracy-min 0.85. To assess performance across varying sequencing depths, datasets were simulated at gradients of 5×, 10×, 15×, 20×, 25×, 30×, 35×, 40×, 45×, and 50× coverage.

Evaluation criteria were defined as follows:

- True Positive (TP): An eccDNA identified by the pipeline that aligns to a simulated circular sequence with >90% identity (lenient criterion) or >99% identity and <10 bp positional deviation (strict criterion) using bedtools intersect and MUMmer3.
- False Positive (FP): An eccDNA identified by the pipeline that does not meet the above criteria.
- False Negative (FN): A simulated circular sequence not detected by the pipeline.
- True Negative (TN): A simulated linear sequence correctly identified as non-eccDNA. Performance metrics were calculated as:
- Precision: It refers to the proportion of samples that are actually positive among those predicted to be positive by the model.

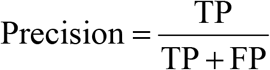

- Recall: It refers to the proportion of samples that are correctly predicted as positive by the model among all actual positive samples.

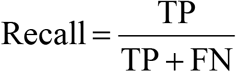

- F1-score: It is a metric that takes both precision and recall into account and balances the two.

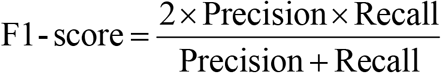

### Bioinformatic Analysis

The sequencing raw data were base-called using Guppy software version 6.3.4, and adaptor/barcode sequences were removed using Porechop to generate clean reads in FASTQ format. Quality control of the FASTQ files was performed using Nanoplot, followed by alignment of the reads to the reference genome GRCh38.p14 using Minimap2. EccDNA identification and result filtering were conducted using ECCFP, eccDNA_RCA_nanopore, CReSIL, and FLED according to their respective guidelines.

The number, length distribution, chromosomal distribution, and breakpoint motifs of eccDNAs were visualized using R. Statistical comparisons between datasets were performed using the Wilcoxon rank-sum test in GraphPad Prism 8.0 (GraphPad Software, San Diego, California, USA). Statistical significance was defined as (P < 0.05), with all probabilities being two-tailed. The overlap of eccDNA results from different analysis pipelines was analyzed using R packages and Bedtools. Consensus sequences generated by different pipelines were compared using MEGA software. Non-parametric tests were applied for variance analysis.

### Verification of eccDNA by PCR and Sanger sequencing

Primers were designed upstream and downstream of the breakpoint sites of the eccDNA for PCR amplification (Supplementary Table S3). Qubit 4.0 was used for quantification, and fragment distribution was detected by the S2 cartridge. The PCR products were tested by Sanger sequencing and the sequencing results were compared with consensus sequence from different bioinfomatical pipelines by MEGE software to show the accuracy of the analysis results.

## Result

### EccDNA identification workflow

ECCFP is an efficient and accurate analysis pipeline specifically designed for identifying eccDNA from enriched long-read sequencing samples. We extracted eccDNAs from the HepG2 cell line as a model and detected them by nanopore sequencing, considering the high performance of the Oxford Nanopore sequencing platform^[25]^. After rigorously removing linear DNA and mitochondrial DNA, we employed rolling circle amplification (RCA) technology to enrich eccDNA and debranched the products by T7 endonuclease^[34]^ (Figure 1A), following the library construction and sequencing. Subsequently, through the pipeline of ECCFP, we demonstrated its working principle and output results.

**Figure 1.**
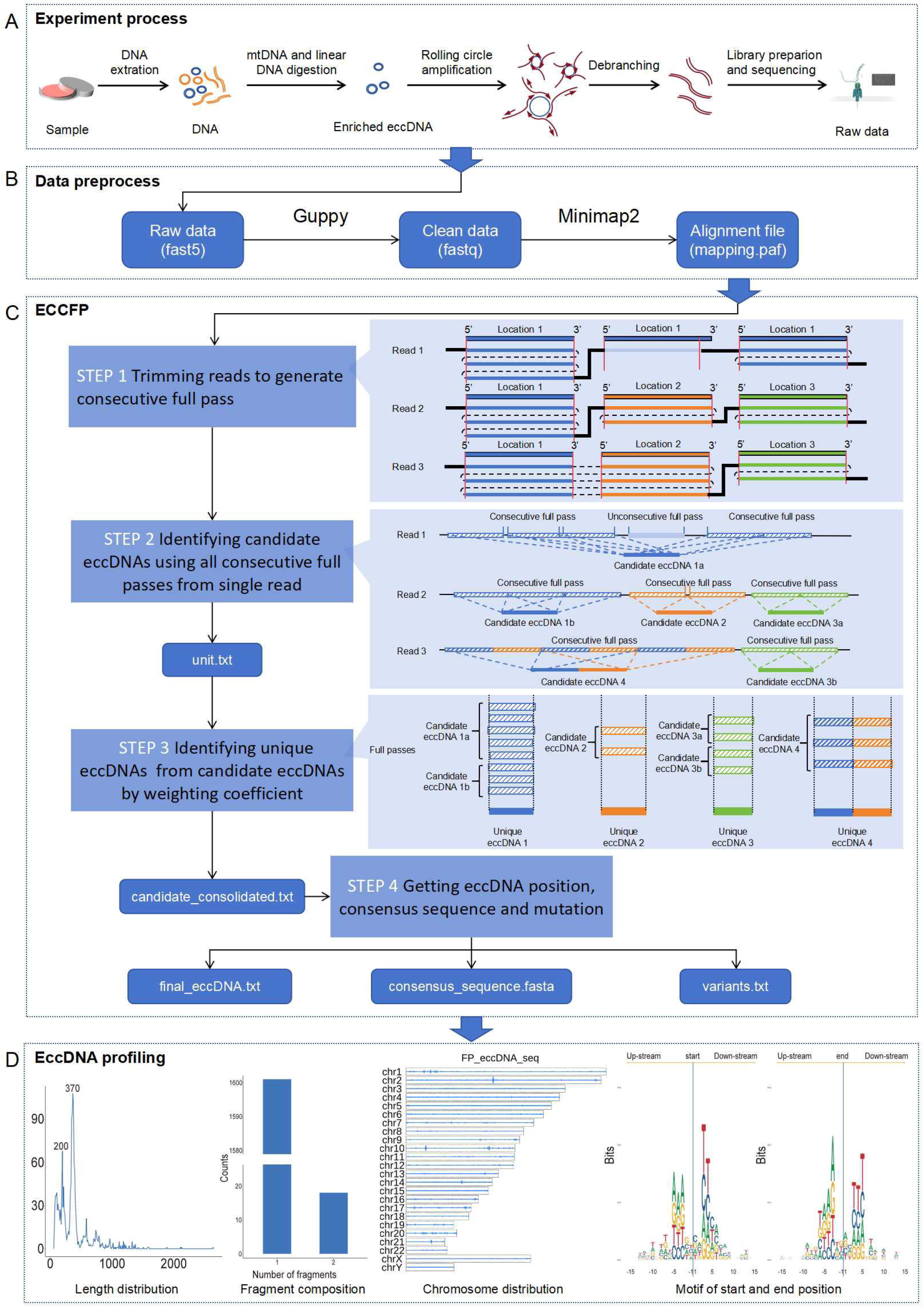
Workflow of eccDNA identification by ECCFP. (A) Experiment process of eccDNA enrichment and nanopore sequencing. (B) Data preprocess using guppy and minimap2. (C) The main pipeline of ECCFP is on the left and the diagrams of common analysis conditions are on the right. STEP 1: Trimming mapping reads based on the alignment file *mapping*.*paf* and preset parameters to generate consecutive full pass. Three common conditions of reads mapping to reference genome are showed in the right diagram. The same color fragments in reads present consecutive full passes from the same location in chromosomes. The light blue fragment is discarded because it does not meet the requirements of consecutive full pass. STEP 2: Using all consecutive full passes from single read to identify candidate eccDNAs. Three common conditions of full passes in reads are showed in the right diagram. In read 1, a single read can identify only one candidate eccDNA. In read 2, a single read can identify more than one candidate eccDNA. In read 3, a single read can identify multi-fragment eccDNAs. STEP 3: Identifying the unique eccDNA position using all consecutive full passes and consolidating the candidate eccDNAs from the same location. Four common conditions of consolidating candidate eccDNAs are showed in the right diagram. STEP 4: Obtaining consensus sequences and identifying mutation by integrating all fragments of consecutive full passes from reads. (D) Routine eccDNA profiling using ECCFP output.

Preprocessing raw data must be performed before initiating the ECCFP workflow (Figure 1B). The fast5 files were converted to fastq format using Guppy, to remove nanopore sequencing adapters and barcode sequences. Subsequently, the data were aligned to the reference genome using Minimap2^[35]^, generating a alignment file (mapping.paf) as the input file.

The ECCFP pipeline consists four major steps to ensure the accuracy of results (Figure 1C). **In step 1**, ECCFP uses the alignment file mapping.paf to trim tandem repeat units with a mapping quality (MapQ) ≥30 from reads and ascertains whether they are consecutive full pass sequences. During trimming, the definition of consecutive full passes is that the tandem repeat sequences on a read align to the same region on the chromosome and the base pair difference between two consecutive full passes on the read is less than 20 bp. Overlapping regions between repeat units are removed to prevent redundant use of overlapping sequences, while gaps between units are preserved. This ensures gaps are not filled using reference genome sequences, thereby avoiding potential artifacts from non-eccDNA sequences. **In step 2**, candidate eccDNAs are identified. When the gap is larger than 20bp, the sequence following the gap is considered a ghost sequence in other analysis pipelines. Consecutive full pass sequences are subsequently identified within the ghost sequence. Ultimately, all consecutive full passes from the same chromosomal location are consolidated into a single candidate eccDNA. Notably, template switching during RCA may result in multiple candidate eccDNAs from a single read. All candidate eccDNAs and their corresponding supporting full pass information are maintained in the intermediate file unit.txt. **In step 3**, unique eccDNAs are obtained. Candidate eccDNAs derived from different reads but at the same chromosomal location are consolidated, requiring that their start and end positions deviate by no more than 20 bp and exhibit consistent alignment orientation. The final junction sites are determined by selecting genomic coordinates supported by the maximum number of supporting full passes. ECCFP improves precise position of eccDNA through this consolidating strategy, significantly reducing false positives. The information of consolidation is maintained in the intermediate file candidate_consolidated.txt. To reflect RCA-derived molecular authenticity, the pipeline retains eccDNAs with divergent orientations at identical position while maintaining fidelity to the original full passes observed in sequencing reads. These orientation-specific results are provided in output file final_eccDNA.txt for user-defined filtration. **In step 4**, a consensus sequence is derived from all consecutive full passes used for identification. These sequences appear as tandem repeats on the reads, originating from the rolling circle amplification of eccDNA. Considering nanopore sequencing errors, a nucleotide is designated as a mutational event only if both the minimum depth of 4× coverage (default) and a 75% frequency threshold (default) are simultaneously satisfied. Otherwise, the pipeline reverts to the reference base at that position. Ultimately, ECCFP generates three output files: final_eccDNA.txt, consensus_sequence.fastq, and variant.txt.

In the data analysis of the HepG2 cell line, ECCFP identified a total of 1,619 unique eccDNA molecules. Their lengths were predominantly concentrated around 200 bp and 370 bp, primarily composed of single fragments, and were more frequently distributed in chromosomal regions of chr1, chr2, and chr10 (Figure 1D). The motifs in the junction sites exhibited a pronounced AT preference and head-to-tail complementarity, consistent with the results of traditional eccDNA studies^[36,37]^.

### Number of candidate eccDNAs identification using all consecutive full passes

During the RCA process, the complexity of reads is incrseased due to amplification errors. We observed five major phenomena (Figure 2A): First, the amplification template of a single read may contain multiple eccDNA molecules (Figure 2A, type 1), implying that a single read can be mapped to multiple chromosomal locations and obtains more than 2 candidate eccDNAs. Second, although some reads contain consecutive full passes originating from the same region, the alignments within these reads are not continuous. In some reads, consecutive full passes are located before the single full pass, while in others, it is positioned after the fragment (Figure 2A, type 2 and 3). Additionally, there are reads where multiple segments of consecutive full passes are discontinuous (Figure 2A, type 4). Existing analysis pipelines typically retain only the longest alignment segment starting from the beginning of the read (the deep blue blocks in Figure 2A), while discarding the other parts as ghost sequences (the blocks filled with slanted line in Figure 2A)^[30]^. In contrast, ECCFP chooses to retain all consecutive full passes that meet the quality criteria, thereby maximizing data utilization while strictly controlling the possibility of erroneous sequences.

**Figure 2.**
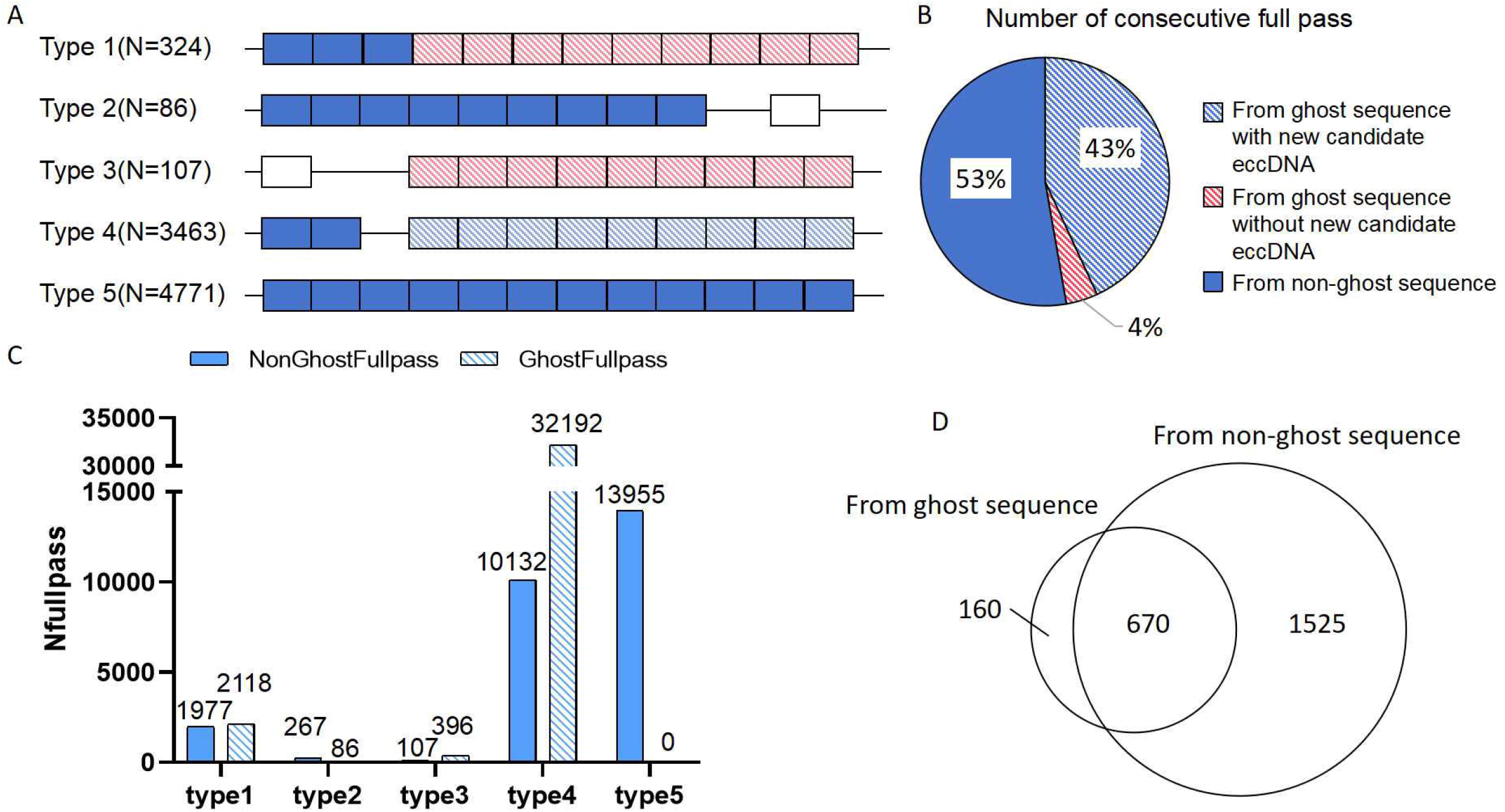
Reads utilization and number of candidate eccDNAs identified by consecutive full passes. (A) Categorization and counts of reads for eccDNA identification. The diagram shows five kinds of full pass conditions in single read. One block presents one full pass. Deep blue blocks present consecutive full passes from non-ghost sequence. Blank blocks mean unconsecutive full passes. Red blocks filled with slanted line present consecutive full passes from ghost sequence and supporting new candidate eccDNA. Blue blocks filled with slanted line present consecutive full passes from ghost sequence and supporting the same candidate eccDNA as deep blue blocks. (B) Pie chart of consecutive full pass from different part of reads. (C) Number of full pass from non-ghost and ghost sequence in different types of reads. (D) Venn diagram showing candidate eccDNAs from two parts: candidate eccDNA identified from non-ghost and ghost sequence.

In the sequencing data of the HepG2 cell line, if ghost sequences are discarded as done by other analysis pipelines, only 53% of full passes are fully utilized (Figure 2B). In contrast, ECCFP, by retaining all consecutive full passes, additionally leverages nearly 45% of reads and uses more 47% of consecutive full passes (Figure 2A, 2B). For example, the read aa769309-4250-4f77-a169-856d3744fd8a is a typical case (Supplementary Table S1). ECCFP is also capable of identifying multiple candidate eccDNAs from a single read, obtaining a total of 2,355 candidate eccDNAs from 8,751 reads. We categorize candidate eccDNAs according to the part of reads. The Venn diagram shows that ECCFP additionally obtained 830 candidate eccDNAs from ghost sequence in reads, among which 160 are newly identified eccDNA molecules (Figure 2D). These findings indicate that, under stringent quality control conditions, retaining ghost sequences aids in the identification of eccDNA.

### Accurate position of eccDNAs identification by consolidating the candidate eccDNAs

To minimize false-positive results, ECCFP employs a consecutive full pass based consolidation strategy during candidate eccDNA integration (Figure 1C, Step 3). Applied to sequencing data from HepG2 cells, this approach consolidates multiple candidate eccDNAs into singular unique eccDNAs (Supplementary Table S2). Candidate eccDNAs can be categorized into two groups: consolidated and non-consolidated (Figure 3A). After consolidating, the number of candidate eccDNAs is reduced from 2,355 to 1,619 unique eccDNAs, with 1,034 candidate eccDNAs being consolidated into 298 unique eccDNAs, which means that ECCPF eliminates 736 potential false-positive results (Figure 3B). This represents a significant reduction of the false-positive rate by 31.2%. The most notable consolidating example is the consolidation of 37 candidate eccDNAs into a single unique eccDNA located at chr5:181126422-181126781(+) (Figure 3C). Leveraging the biological principle that each RCA event generates a full pass corresponding to the template eccDNA, the algorithm utilizes full pass count as a consolidating parameter. The start and end positions of all consecutive full passes used to identify this eccDNA are concentrated at chr5:181126422 and chr5:181126781, accurately reflecting the eccDNA position.

**Figure 3.**
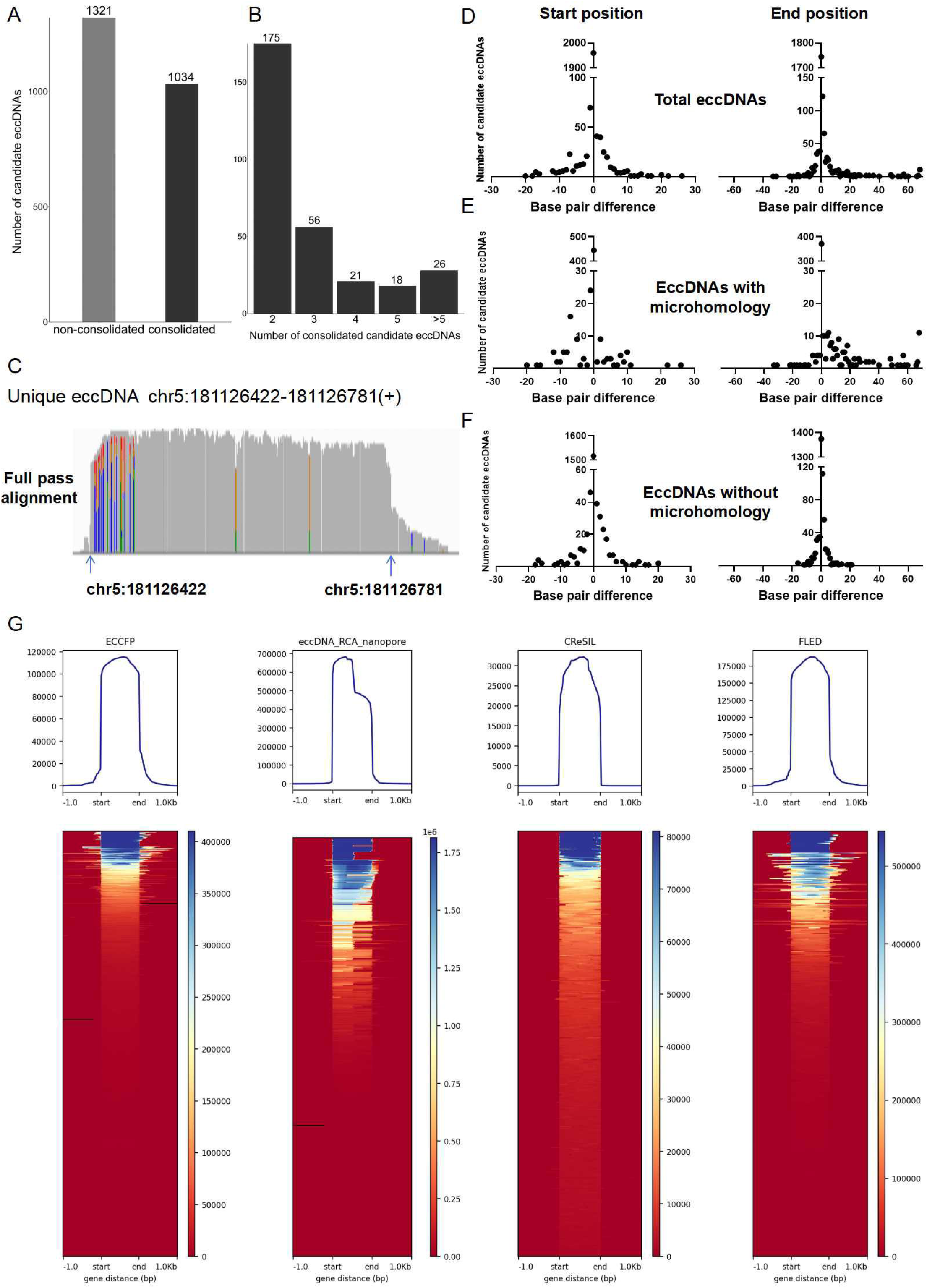
Unique eccDNAs identification by weighting all consecutive full passes and consolidating candidate eccDNAs. (A) Number of candidate eccDNAs categorized as consolidated and unconsolidated. (B) Number of unique eccDNAs with varying numbers of consolidated candidate eccDNAs. (C) Example of full-pass alignment for unique eccDNA on chr5:181126422-181126781(+). (D) Number of candidate eccDNAs corresponding to different base pair differences between total unique and candidate eccDNAs. (E) Number of candidate eccDNAs corresponding to different base pair differences between unique and candidate eccDNAs with junction flanking sequences containing microhomology. (F) Number of candidate eccDNAs corresponding to different base pair differences between unique and candidate eccDNAs with junction flanking sequences lacking microhomology. (G) Profile and heatmap plots present the base pair difference between alignments and unique eccDNAs from four analysis pipelines.

The base pair difference between the start and end positions of unique eccDNA and their corresponding candidate eccDNA indicates that ECCFP is capable of obtaining the positions of eccDNA as accurately as possible (Figure 3D). The start and end positions of most consecutive full passes accurately reflect the position of the template eccDNA, while a minority show significant base pair difference. Considering the impact of microhomology sequence on junction site, candidate eccDNA is categorized into two groups (Figure 3E-F). Comparison of the base pair difference between candidate eccDNAs and unique eccDNAs, the results show that candidate eccDNA with microhomology has greater end position deviations, but ECCFP achieves the effective redundancy of the false-positive results.

We also compared the position of unique eccDNAs from analysis pipelines with actual genomic position of sequencing data (Figure 3G). ECCFP obviously detected the start and end positions of alignment in reads as possible. FLED performs inferior in this respect. There is significant difference between unique eccDNAs identified by eccDNA_RCA_nanopore and alignments, because eccDNA_RCA_nanopore uses reference genome sequence to fill the gap between full passes leading to this bias. The results of CReSIL show that unique eccDNA regions are oversized, almost completely including the actual genomic region of the sequencing data. The above situation demonstrates that eccDNA position identified by ECCFP is more correspond to the actual genomic region of the sequencing data, which presents the real eccDNA on the genome.

### Performance of ECCFP comparing with other pipelines

We generated two simulated datasets (see Methods) to comprehensively evaluate the performance of ECCFP and compared it with existing analysis pipelines (such as eccDNA_RCA_nanopore, CReSIL, and FLED). When setting up the simulated eccDNA data, we randomly selected 5,000 regions from the reference genome as positive sequences (circular DNA) and 5,000 regions as negative sequences (linear DNA), ensuring a consistent length distribution between them (Figure 4A), and subsequently generated long-read sequencing data for both types of sequences. Simulated data without ghost sequence generated according to the references did not produce ghost sequences containing multiple candidate eccDNA, whereas our designed simulated data according to the proposion of eccDNA sequencing data from HepG2 cells better reflect actual sequencing scenarios (Figure 4B). After implementing stricter criteria (see Methods), the evaluation results for simulated data including ghost sequences showed significant changes: ECCFP demonstrates excellent performance, with significantly higher precision and recall compared to other analysis pipelines (Figure 4CD). Its F1-score remains stable above 0.9 across varying sequencing depths, and other pipelines show more pronounced sensitivity to sequencing depth fluctuations (Figure 4E). Specifically, eccDNA_RCA_nanopore exhibits unstable precision below 0.7 that decreases with increased sequencing depth, significantly impacting its F1-score which consequently declines. Although its recall remains high and correlates positively with sequencing depth, this comes at the cost of substantially elevated false positives (Figure 4G). CReSIL shows sub-optimal performance across all metrics, with progressive deterioration as sequencing depth increases (Figure 4H). FLED maintains moderate precision but suffers from low recall, displaying similar trends to ECCFP albeit with consistently inferior numerical values (Figure 4I). Crucially, ECCFP consistently outperformed other pipelines in precision and recall across all coverage depths, with statistically significant advantages. It demonstrated superior F1-scores, enhanced stability, and better comprehensive performance when analyzing low-depth sequencing data. Moreover, ECCFP achieved better comprehensive scores when analyzing low-depth sequencing data. These findings highlight ECCFP’s broader applicability and robustness. Given the current prevalence of moderate-depth eccDNA sequencing data, ECCFP effectively fulfills most eccDNA detection requirements for long-read sequencing. Notably, ECCFP shows particular strength in identifying small eccDNA fragments and outperforms competing pipelines when handling low-depth sequencing data.

**Figure 4.**
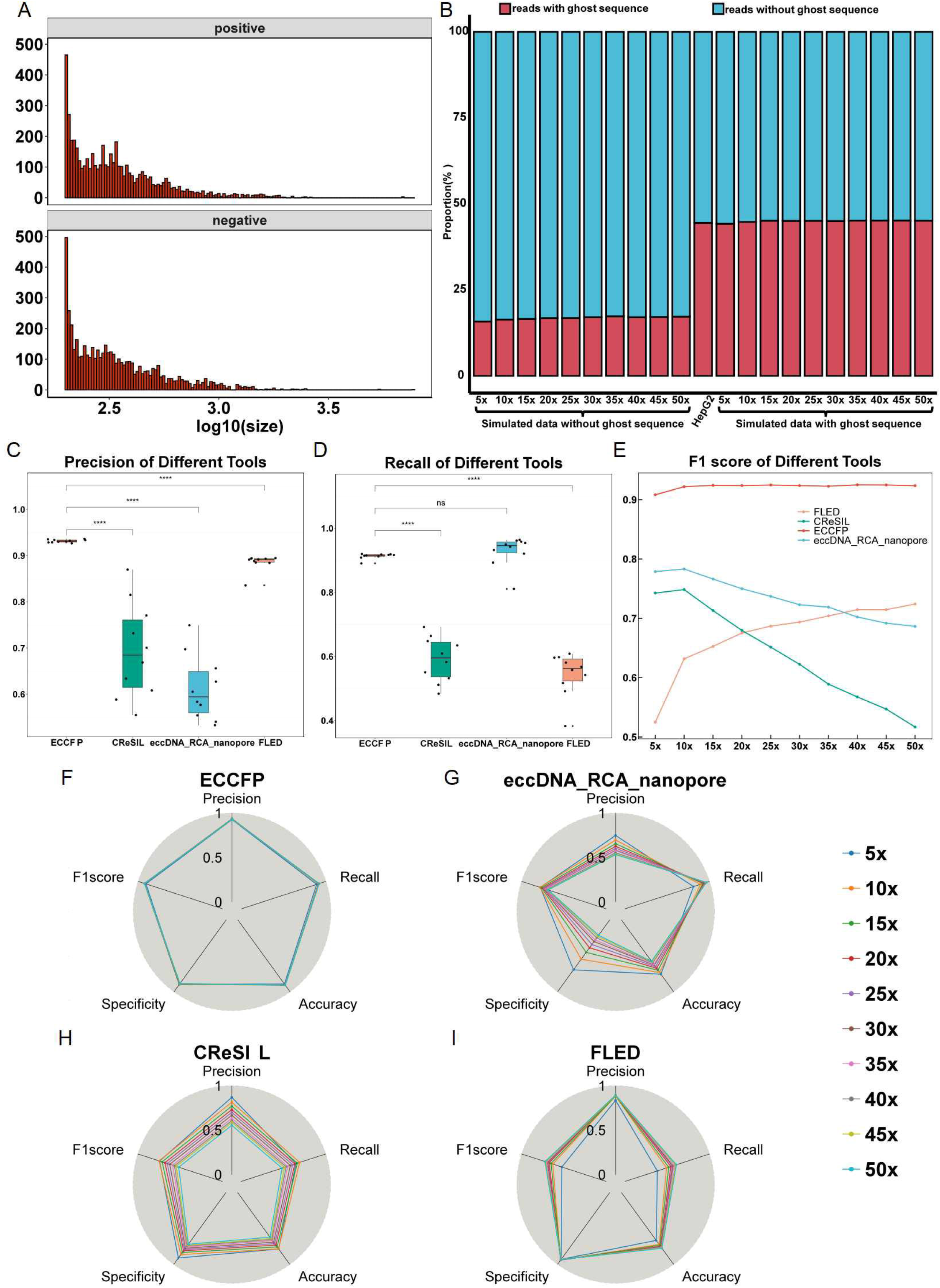
Evaluating ECCFP using simulated data. (A) Size distribution of the simulated eccDNA positive and negative datasets. (B) Proportion of reads with ghost sequences in three datasets: simulated data without ghost sequence, nanopore sequencing of eccDNAs from HepG2 cell line and simulated data with ghost sequence. (C) Precision of analysis pipelines for simulated data including ghost sequences under strict criterion. (D) Recall of analysis pipelines for simulated data including ghost sequences under strict criterion. (E) F1-score of analysis pipelines for simulated data including ghost sequences under strict criterion. (F-I) Evaluation of four analysis pipelines using simulated data including ghost sequences at varying sequencing depths.

In the analysis of real sequencing data, ECCFP demonstrated equally strong performance. In HepG2 cell line results, eccDNA_RCA_nanopore detected 2,105 eccDNAs—substantially more than other methods. ECCFP detected 1,619 eccDNA molecules, including molecules from different strands at the same location. CReSIL and FLED detected 448 and 446 eccDNAs, respectively. When requiring complete positional identity across all four methods, ECCFP showed the greatest overlap with other pipelines (Figure 5A). Relaxing the criteria to 90% sequence length overlap using bedtools further revealed that ECCFP encompassed 70-80% of results from other methods (Figure 5B). Chromosomal distribution also indicated that ECCFP results evenly encompassed results from other analysis pipelines (Figure 5C). These same locations suggest that the start and end positions identified by these pipelines may not always be accurate. To validate the hypotheses regarding the results, five unique eccDNA molecules were selected for PCR amplification and Sanger sequencing (Supplementary Figure 1). Consensus sequences most closely matching the Sanger results were chosen from each analysis method and compared using MEGA software (Table 1). As shown in the figures, ECCFP output sequences exhibited strong concordance with experimental results, while CReSIL and FLED results displayed deviations in start and end positions and sequence content (Figure 5D). These findings demonstrate that ECCFP not only identifies more eccDNA molecules but also significantly reduces false positives while maintaining high accuracy and reliability.

**Table 1.**
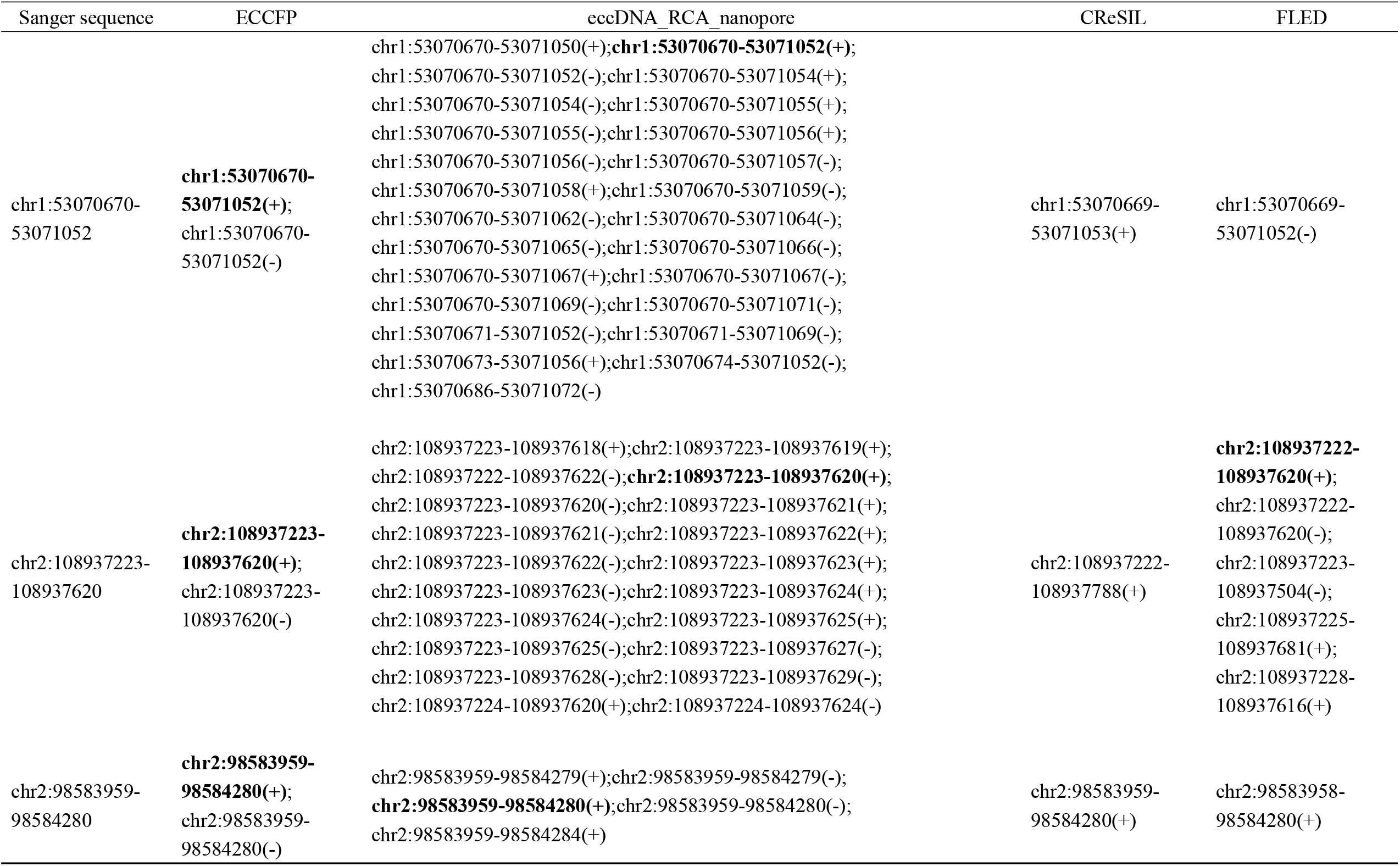
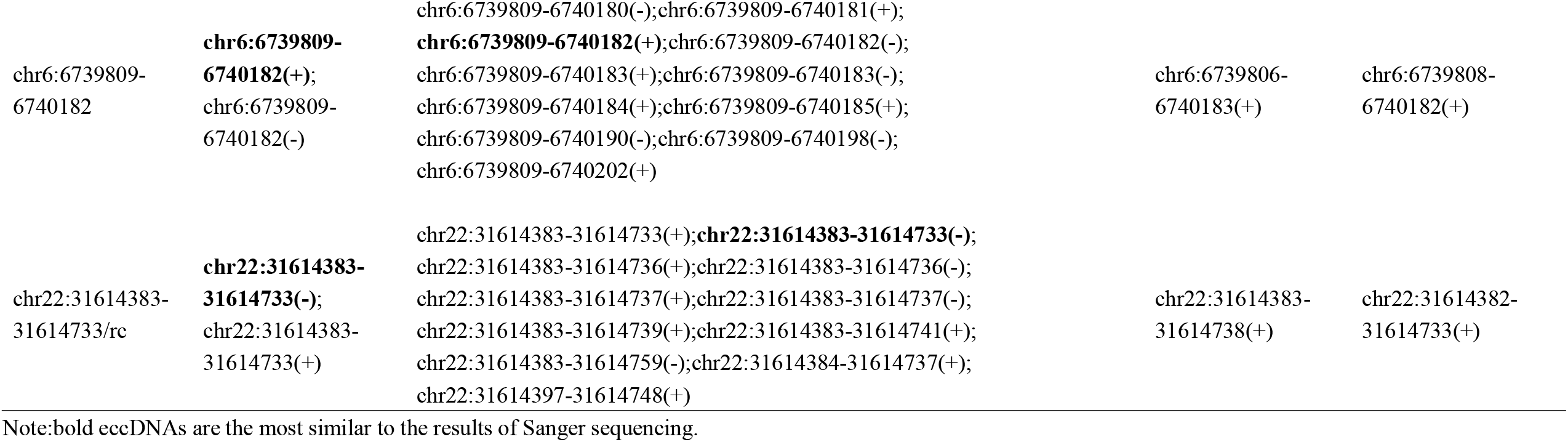
Chromosome position identitied by different eccDNA analysis pipelines and validated by Sanger sequencing

**Figure 5.**
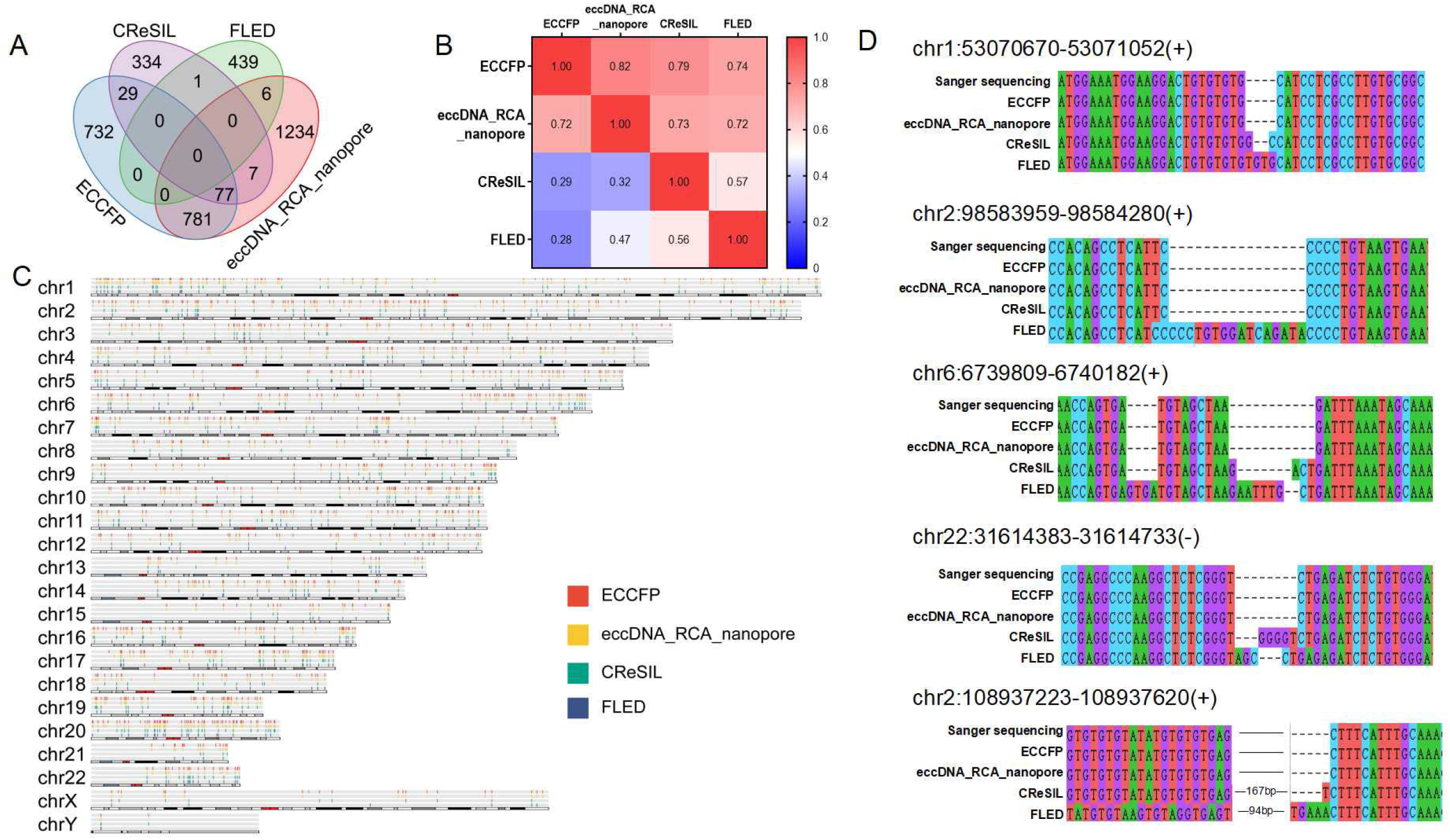
The correlation and junction sequence validation for current eccDNA analysis pipelines. Comparison of the results of the four analysis pipelines on eccDNA sequencing from HepG2 cell lines. (A) Venn diagram showing eccDNA numbers identified by four analysis pipelines. (B) 90% overlap of eccDNA positions among the four results. (C) Chromosomal distribution of eccDNAs identified by four tools. (D) Sanger sequencing results aligned with consensus sequences of five verified eccDNAs generated by four analysis pipelines.

## Discussion

ECCFP is a novel analysis pipeline that significantly reduces the false positive rate while maintaining high sensitivity and accuracy. By integrating both read-based and fragment-based analysis pipelines, it optimizes the utilization of ghost sequences traditionally discarded in conventional pipelines. This pipeline retains all consecutive full passes on reads and employs full pass as a weighting parameter to consolidate candidate eccDNAs, thereby enhancing the reliability of the analysis.

ECCFP can identify more candidate eccDNAs because it retains all consecutive full passes on reads, including ghost sequences discarded by other analysis pipelines. Under ideal conditions, the products of RCA are characterized by tandem repeat units observed in sequencing reads^[38,39]^. For long-read sequencing data, the widely accepted identification principle involves locating tandem repeat sequences from reads that can be aligned to the reference genome, which are recognized as eccDNA sequences^[40]^. Different pipelines employ various methods to infer the chromosomal origin and sequence of the template eccDNA based on these tandem repeat units. However, the complexity of reads increases because of amplification errors during RCA. Since phi29 DNA polymerase may switch between DNA strands during RCA^[41]^, generating substantial inverted repeat units, these produces are typically labeled as ghost sequences and discarded by other pipelines. In this study, we demonstrate that these sequences filterd with strict requires can actually be effectively utilized to discover new candidate eccDNA sequences, maximizing the use of these data and thereby significantly improving the efficiency of eccDNA detection.

ECCFP enables more accurate identification of eccDNAs. Existing eccDNA analysis pipelines have certain limitations. For example, while eccDNA_RCA_nanopore demonstrates the capacity to detect abundant eccDNA molecules, it generates a high proportion of false-positive results. CReSIL and FLED need improvement in the accuracy of start and end positions as well as the stability of consensus sequences (Figure 5D). In contrast, ECCFP employs a full pass-based consensus consolidating strategy that significantly reduces false-positive outcomes while ensuring comprehensive identification of eccDNA molecules. Furthermore, ECCFP exhibits enhanced capability in detecting small eccDNAs, which are typically underrepresented due to the preferential amplification of smaller circular templates during RCA processes^[42]^.

To more rigorously evaluate the performance of existing analysis pipelines, we developed simulated datasets incorporating ghost sequences to reflect complexities encountered in actual sequencing scenarios. Since eccDNA molecules predominantly range between 200-400 bp in size, previous studies employed overly lenient criteria for defining true positives (TPs)^[40,43]^, leading to inflated TP counts (Supplementary Table S4). Previous studies employed relatively lenient criteria for true positive (TP) identification, resulting in overestimated TP counts that compromised accurate assessment of precision and recall rates (Supplementary Figure 2-5). Our analysis demonstrates ECCFP’s consistent superior performance, as evidenced by its higher F1-scores across varying sequencing depths compared to other pipelines. Notably, significant differences were observed when evaluating analysis pipelines using simulated data without ghost sequence versus ghost sequence-containing data (Supplementary figure 6 and Figure 4C-E): ECCFP and CReSIL demonstrated robust and stable performance. FLED showed improved precision but a substantial decline in recall rate. The precision of eccDNA_RCA_nanopore slightly decreased. In simulated data with ghost sequence under strict criteria, eccDNA_RCA_nanopore exhibited substantial precision deterioration with increasing sequencing depth due to its fundamental reliance on single-read identification for eccDNA, leading to corresponding false-positive rate increases and F1-score declines. CReSIL showed marked reductions in both precision and recall under strict criteria, particularly pronounced at higher sequencing depths (Figure 4H). This phenomenon likely stems from its consensus consolidating approach that incorporates discontinuous full passes, generating progressively elongated eccDNA sequences that increasingly deviate from simulated positive sequences. Although FLED maintained relatively high precision, its limited eccDNA detection capacity may be attributed to discarding ghost sequences and stringent alignment filtering criteria, unlike ECCFP’s pipeline that effectively utilizes consecutive full passes on ghost sequences.

In the evaluation of actual sequencing data, the advantages of ECCFP become even more pronounced. Compared with other analysis pipelines, ECCFP not only identifies more eccDNA molecules but also significantly reduces false positives. Through PCR amplification and Sanger sequencing validation, the consensus sequences output by ECCFP are highly consistent with validated results. In contrast, other pipelines (such as CReSIL and FLED) exhibited varying degrees of discrepancies in the positions and sequence content. These results further demonstrate the superiority of ECCFP in terms of accuracy and reliability.

ECCFP, as a novel analysis pipeline, is specifically designed to identify eccDNA from long-read sequencing data enriched through RCA. This pipeline combines the advantages of existing analysis pipelines and is optimized for the characteristics of reads processed by RCA. Regrettably, ECCFP is limited by the propensity of RCA in eccDNA enrichment experiments to preferentially amplify eccDNA molecules smaller than 10 kb^[42]^. It is more suitable for detecting smaller eccDNA molecules, such as microDNA and spcDNA, rather than ecDNA. If future research be able to more effectively address the shortcomings in the experimental aspects, the amplification scope of ECCFP would be significantly broadened. The development of ECCFP provides new insights for optimizing other long-read sequencing-based analytical methods, advancing the application of sequencing technologies in complex genomic region analysis. In the future, ECCFP is expected to play an important role in both basic research and clinical translation, facilitating the transition of eccDNA research from mechanistic exploration to practical application.

## Supplementary data

Supplementary data are available online.

## Author contribution

This study was conceptualized and designed by S.W. The experiment and Nanopore sequencing were performed by T.Z. , M.L. , B.M. and J.L. The pipeline development and data analysis were completed by W.L. , T.Z. , Q.Z. , J.Z. and S.C. The manuscript was written by T.Z. S.W. and W.L. . The proofreading of manuscripts was completed by S.W. and T.L. All authors read and approved the submission and publication.

## Data availability

The code and usage details of ECCFP pipeline can be found at the website https://github.com/WSG-Lab/ECCFP. All sequencing data has been deposited at National Geno mics Data Center, and available upon request.

