## Supplement Figure for "ECCFP: a consecutive full pass based bioinformatic analysis for eccDNA identification using Nanopore sequencing data"

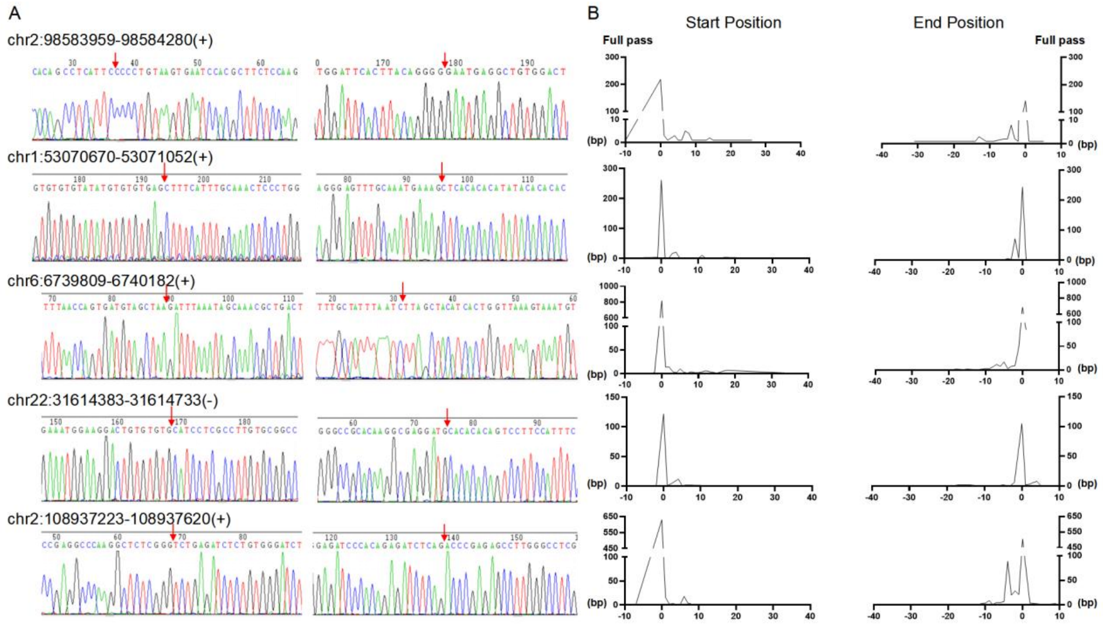


**Supplementary figure 1.** Verified eccDNAs condition. (A) Partial Sanger sequencing results. Left: Forward sequencing result. Right: Reverse sequencing result. The red arrow indicates the eccDNA junction site. (B) Base pair differences between accurate eccDNA and corresponding full passes in the start and end positions of eccDNA.

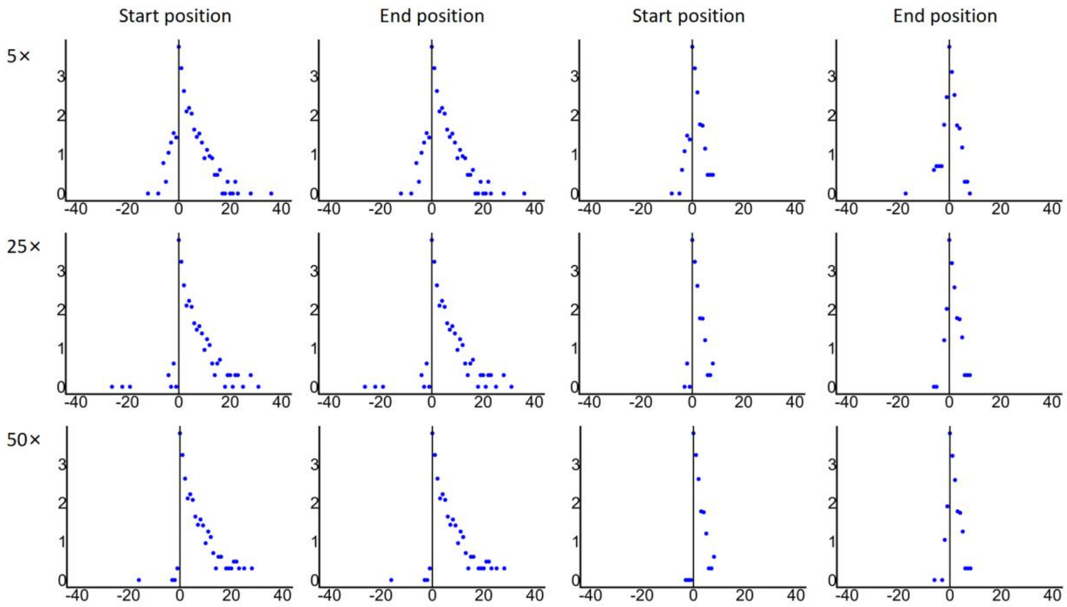


**Supplementary figure 2.** Base pair differences in the start and end position between TP results from ECCFP and true eccDNA of simulated data under different filtering conditions of lenient criterion in the left and strict criterion in the right.

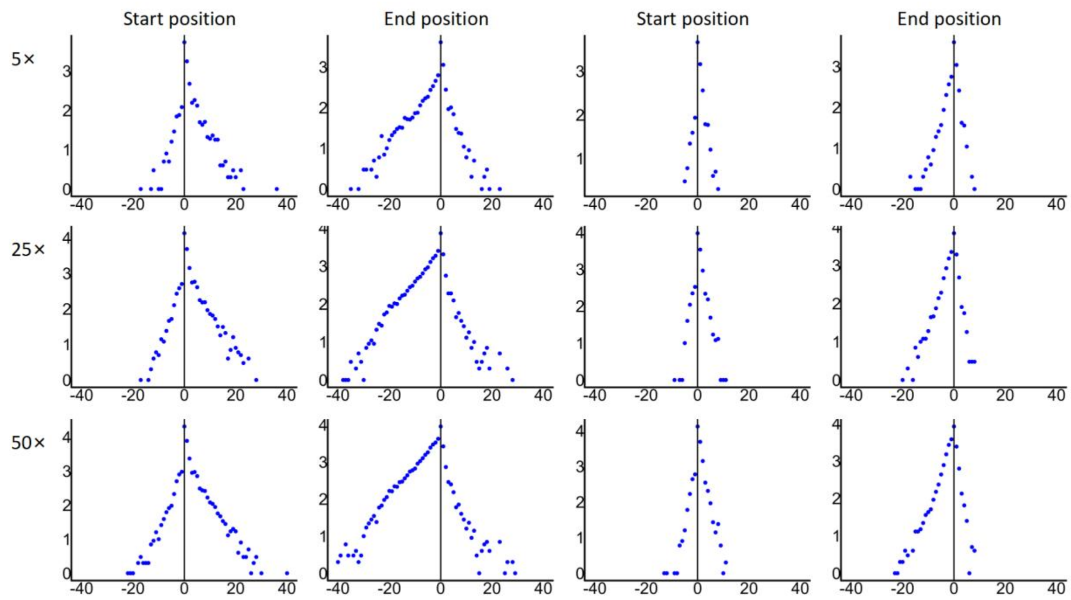


**Supplementary figure 3.** Base pair differences in the start and end position between TP results from eccDNA_RCA_nanopore and true eccDNA of simulated data under different filtering conditions of lenient criterion in the left and strict criterion in the right.

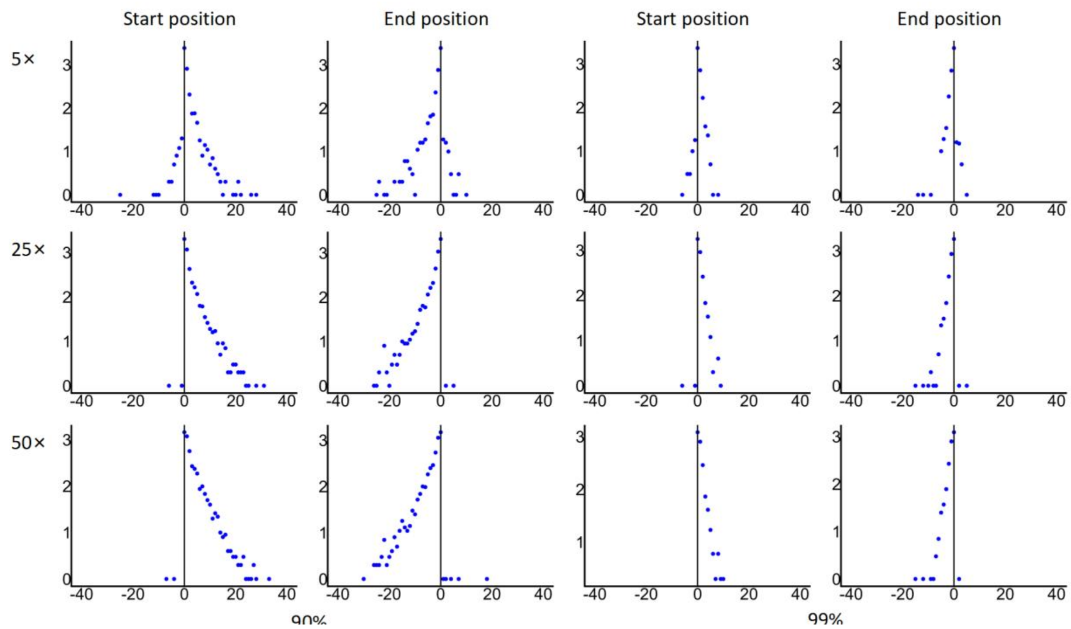


**Supplementary figure 4.** Base pair differences in the start and end position between TP results from CReSIL and true eccDNA of simulated data under different filtering conditions of lenient criterion in the left and strict criterion in the right.

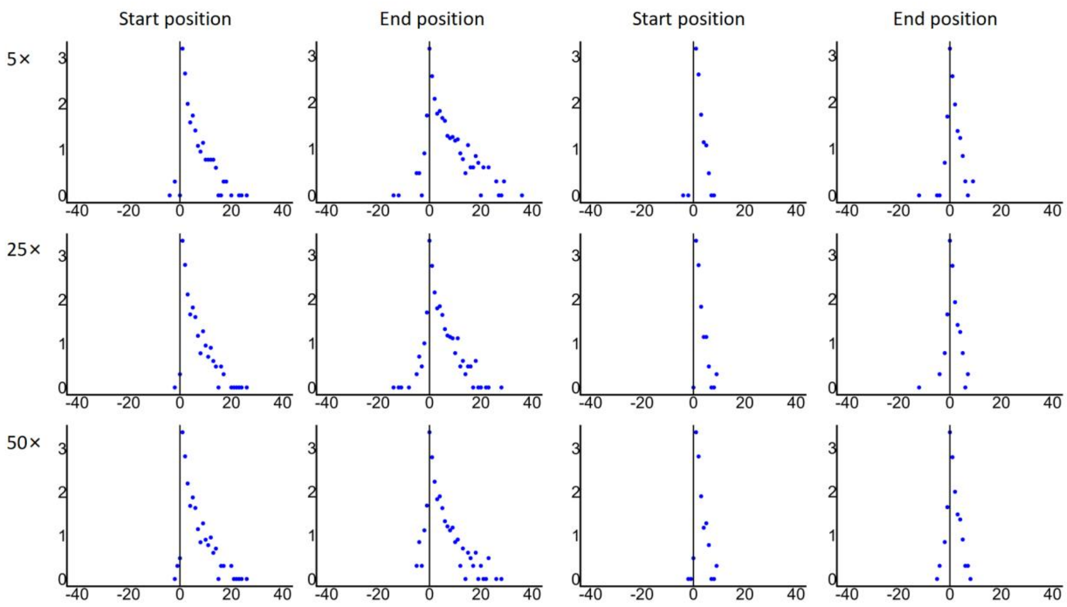


**Supplementary figure 5.** Base pair differences in the start and end position between TP results from FLED and true eccDNA of simulated data under different filtering conditions of lenient criterion in the left and strict criterion in the right.

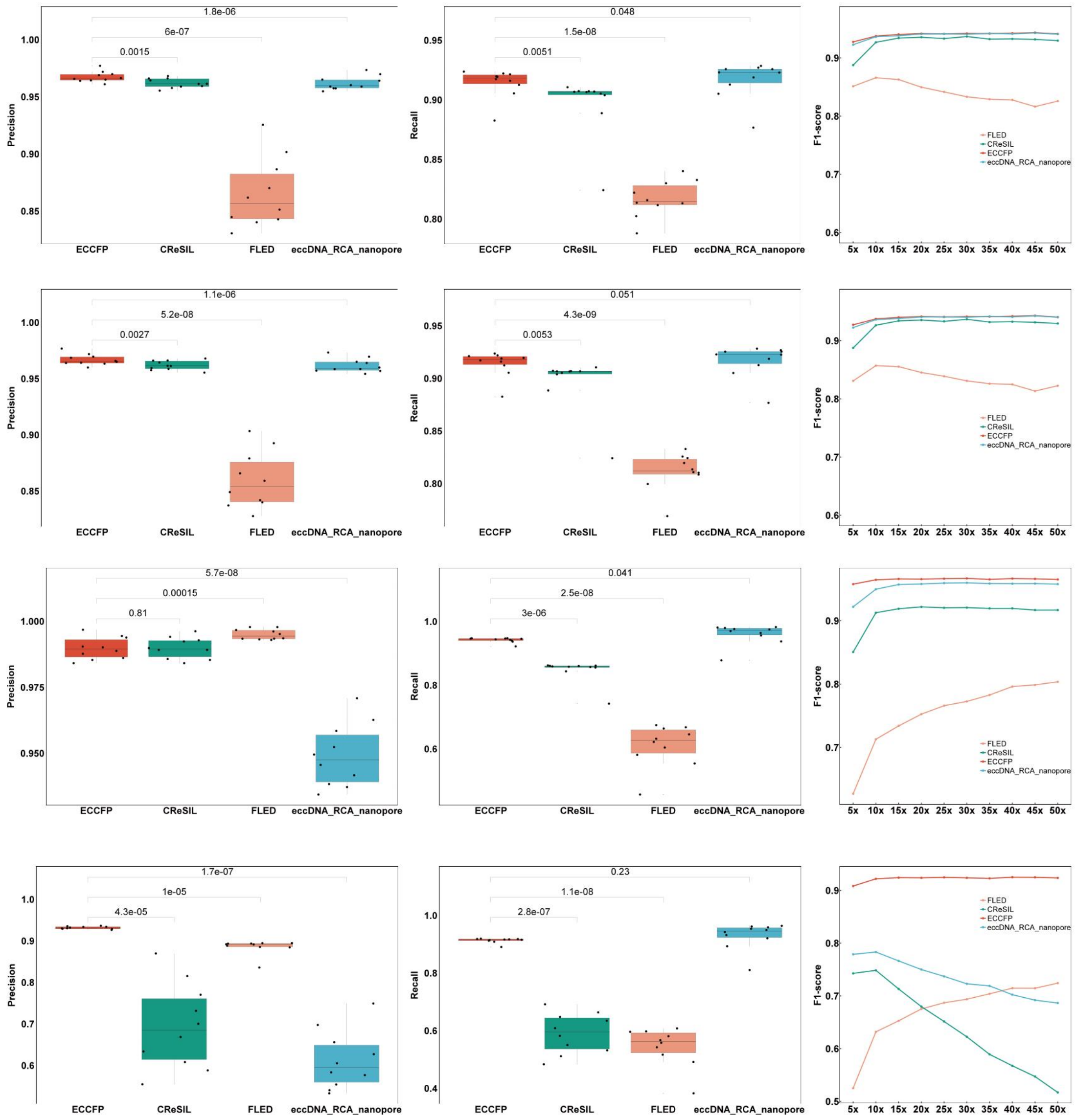


**Supplementary figure 6.** Evaluation of analysis tools under different TP conditions for two types of simulated data. (A) Precision of analysis tools for general simulated data under lenient criterion. (B) Recall of analysis tools for general simulated data under lenient criterion. (C) F1-score of analysis tools for general simulated data under lenient criterion. (D) Precision of analysis tools for general simulated data under strict criterion. (E) Recall of analysis tools for general simulated data under strict criterion. (F) F1-score of analysis tools for general simulated data under strict criterion. (G) Precision of analysis tools for simulated data including ghost sequences under lenient criterion. (H) Recall of analysis tools for simulated data including ghost sequences under lenient criterion. (I) F1-score of analysis tools for simulated data including ghost sequences under lenient criterion. (J) Precision of analysis tools for simulated data including ghost sequences under strict criterion. (K) Recall of analysis tools for simulated data including ghost sequences under strict criterion. (L) F1-score of analysis tools for simulated data including ghost sequences under strict criterion.
